# Epigenetic suppression of interferon lambda receptor expression leads to enhanced HuNoV replication *in vitro*

**DOI:** 10.1101/523282

**Authors:** Sabastine E. Arthur, Frédéric Sorgeloos, Myra Hosmillo, Ian G. Goodfellow

## Abstract

Human norovirus (HuNoV) is the main cause of gastroenteritis worldwide yet no therapeutics are currently available. Here, we utilize a human norovirus replicon in human gastric tumor (HGT) cells to identify host factors involved in promoting or inhibiting HuNoV replication. We observed that an IFN-cured population of replicon-harboring HGT cells (HGT-cured) was enhanced in their ability to replicate transfected HuNoV RNA compared to parental HGT cells, suggesting that differential gene expression in HGT-cured cells created an environment favouring norovirus replication. Microarray analysis was used to identify genes differentially regulated in HGT-NV and HGT-cured compared to parental HGT cells. We found that the IFN lambda receptor alpha (IFNLR1) expression was highly reduced in HGT-NV and HGT-cured cells. All three cell lines responded to exogenous IFN-β by inducing interferon stimulated genes (ISGs), however, HGT-NV and HGT-cured failed to respond to exogenous IFN-λ. Inhibition of DNA methyltransferase activity with 5-aza-2’-deoxycytidine partially reactivated IFNLR1 expression in HGT-NV and IFN-cured cells suggesting that host adaptation occurred via epigenetic reprogramming. In line with this, ectopic expression of the IFN-λ receptor alpha rescued HGT-NV and HGT-cured cells response to IFN-λ. We conclude that type III IFN is important in inhibiting HuNoV replication in vitro and that the loss of IFNLR1 enhances replication of HuNoV. This study unravels for the first time epigenetic reprogramming of the interferon lambda receptor as a new mechanism of cellular adaptation during long-term RNA virus replication and shows that an endogenous level of interferon lambda signalling is able to control human norovirus replication.

**Importance:** Noroviruses are one of the most wide-spread causes of gastroenteritis yet we have no therapeutics for their control and we do not fully understand what cellular processes control viral replication. Recent work has highlighted the importance of type III interferon (IFN) responses in the restriction of viruses that infect the intestine. Here we analysed the adaptive changes required to support long term replication of noroviruses in cell culture and found that the receptor for type III IFN is decreased in its expression. We confirmed that this decreased expression was driven by epigenetic modifications and that cells lacking the type III IFN receptor are more permissive for norovirus replication. This work provides new insights into key host-virus interactions required for the control noroviruses and opens potential novel avenues for their therapeutic control.

## Introduction

With the introduction of the rotavirus vaccine, human norovirus (HuNoV) is now the main etiologic agent responsible for gastroenteritis worldwide (1–4). The disease, characterized by diarrhoea, nausea and vomiting is generally self-limiting in healthy adults. However, in immunocompromised, elderly patients and young children under the age of five, the disease can become chronic and sometimes lead to death mostly due to dehydration (5–9). Despite this and the profound economic burden of the disease (10), there are no therapeutics or licensed vaccines available.

Until the advent of the B lymphocyte-based and stem cell-derived human intestinal enteroid culture systems (11, 12), HuNoV research has been hampered by the lack of cell culture and small animal models recapitulating noroviral infection and pathogenesis. However, limitations in robustness, cost and labour intensity associated with both methods foreshadow that advances in HuNoV research still relies on the well-established murine norovirus (MNV) infection model and norovirus replicon systems (13, 14). These systems have been used to identify key host factors that impact the lifecycle of HuNoV, reviewed by Thorne and Goodfellow (15).

Contrary to the type I IFNs (IFN-α/β) which were discovered more than an half century ago (16), type III interferons (IFN-λ) were discovered only little over a decade (17, 18). Although both cytokine families possess similar functions, a few but crucial differences exist in their biology. Notably, both cytokines signal through distinct heterodimeric receptors with type I IFN signalling through the interferon alpha/beta receptor (IFNAR) composed of the IFNAR1 and IFNAR2 subunits and type III IFN through the interferon lambda receptor (IFNLR) which consists of the interferon lambda receptor 1 (IFNLR1) and interleukin 10 receptor beta (IL10RB) subunits. Unlike the type I IFN receptor that is expressed on most cell types, expression of the IFNLR1 subunit is restricted to cells of the mucosal epithelium, neutrophils and human hepatocytes (19–21). Although the immune cells of the blood were also shown to express the IFNLR1 subunit, the receptor has been reported to lack the ability to respond to IFN-λ (22).

Expression of type I and III interferons is regulated at the transcriptional level and relies on the recognition of conserved pathogen-associated molecular patterns (PAMPs). Detection of these molecular signatures by extracellular and intracellular pattern recognition receptors triggers the coordinated activation of distinct signalling pathways responsible for the activation of IRF-3/7 and NF-κB transcription factors that are required for IFN gene transcription. Secreted IFN-λ then binds to the IFNLR and is thought to activate the Janus kinase 1 and tyrosine kinase 2. Subsequently, recruited signal transducer and activator of transcription 1 (STAT1) and STAT2 are activated through phosphorylation leading to the expression of IFN stimulated genes (ISGs), some of which have direct antiviral activities (23). Interferon lambdas are important players in both innate and adaptive immunity and have profound antiviral effects on a variety of viruses (24–28). Expression of IFNLR1 on epithelial cells of the small intestine and colon was shown to be important in IFN-λ-mediated antiviral activity against persistent MNV and reovirus infection *in vivo* (29). Treatment of persistently infected mice lacking adaptive immune response (Rag1^−/−^) with IFN-λ abolished viral replication, suggesting that IFN-λ can cure persistent MNV infection in absence of adaptive immunity and this ability requires the expression of IFNLR1 (30). In line with this, the effect of antibiotics that inhibit persistent MNV infection in the gut has also been shown to be dependent on IFNLR1 expression as well as IRF3 and STAT1 transcription factors (31). It was observed that AG129 sentinel mice lacking the ability to respond to both IFN-α/β and IFN-γ housed together with MNV-infected mice developed a diarrhoea-associated MNV infection. Overexpression of IFN-λ in sentinel mice upregulated ISG expression, inhibited MNV replication in the small intestine and prevented them from being infected when co-housed with MNV-infected mice (32). This article was submitted to an online preprint archive (33).

## Results

### Generation and characterization of human cell lines bearing stable human norovirus replicons

To understand the influence of viral and host factors involved in HuNoV replication, we sought to generate several human cell lines stably replicating HuNoV RNA. To this end, BHK-21 cells were transfected with capped Norwalk replicon RNA harbouring a neomycin selection marker (14) and subjected to G418 selection 48 hours after transfection (Fig. 1A). Although the vast majority of the cells died within one week, individual cell colonies were observed and subjected to limiting dilution. A single high-expressing clone was selected and expanded to generate stable replicon-containing BHK-21 cells (BHK-NV). VPg-linked RNA extracted from these cells was transfected into HGT cells, a cell line of epithelial origin which was subsequently selected on the basis of their G418 resistance in order to generate human norovirus replicon cells (HGT-NV). These HGT-NV cells were either collected as a population or subjected to limiting dilution to produce HGT-NV cell clones. The HGT-NV population was further passaged 16 times in the presence of IFN□α at a concentration of 1000 U/mL in the absence of G418 selection over an 8 weeks period leading to the generation of HGT-Cured cells. These cells were subsequently cultured in presence of G418 to observe their loss of resistance to G418 confirming the complete elimination of the replicon. Detection of HuNoV RNA by RT-qPCR analysis confirmed the presence of noroviral genomes in HGT-NV cells that were absent from HGT-Cured or parental HGT cells used as control (Fig. 1B). To confirm the presence of authentic steady-state replication of Norwalk RNA, cells were subjected to immunofluorescence analysis using monoclonal antibodies directed against dsRNA, a by-product assumed to be universally generated during viral replication (34, 35). As shown in Fig. 1C, punctate structures reminiscent of replication complexes were identified in HGT-NV cells while no signal above background levels was detected in HGT-Cured or in parental cell lines. Taken together these results suggest that HuNoV VPg-linked RNA successfully replicates in HGT cells and displays all the characteristics of a replication-competent RNA.

**Figure 1.**
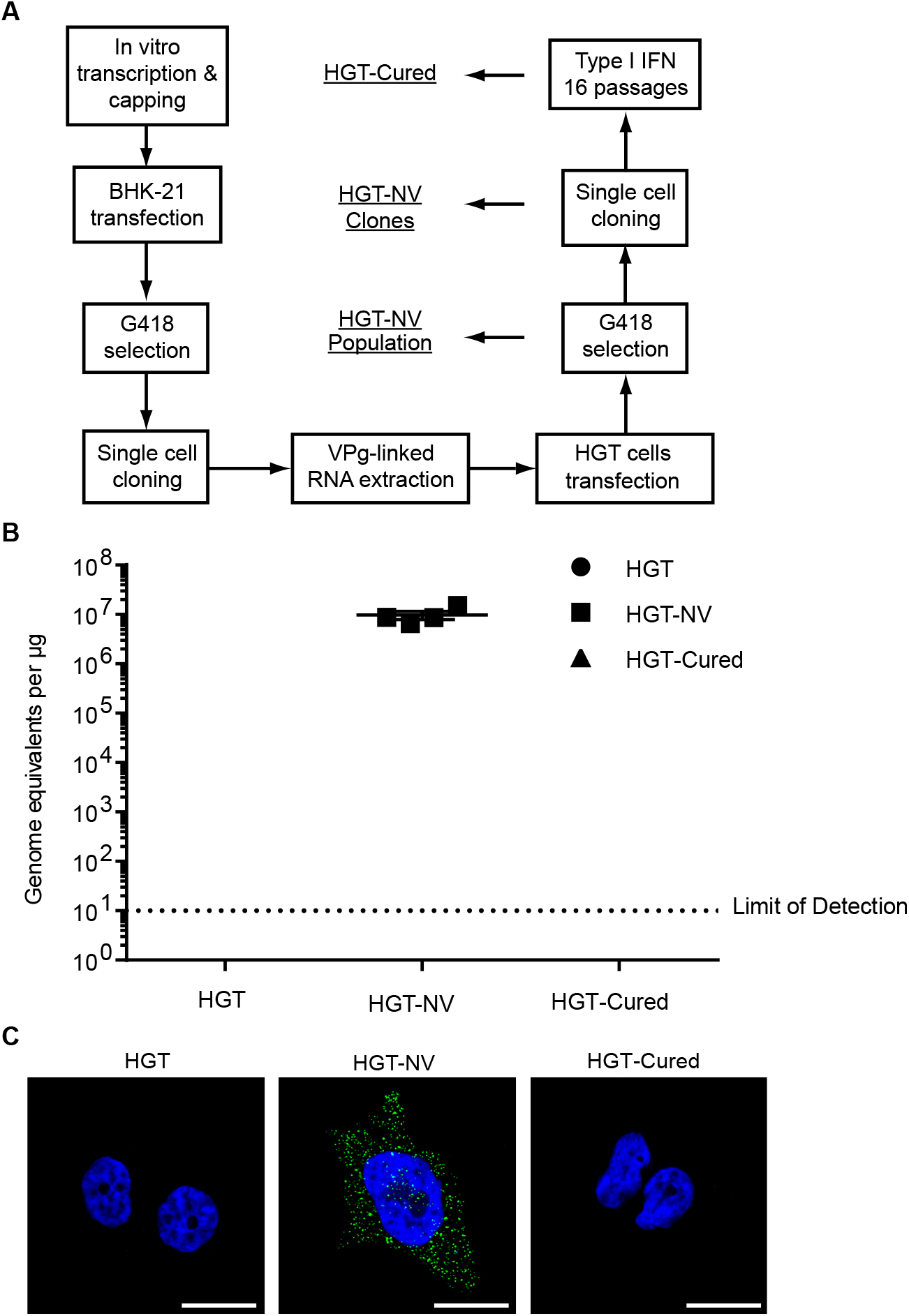
Generation and characterization of stable HuNoV replicons in HGT and U2OS cell lines. A. Diagram showing the steps used to generate the various cell lines carrying human norovirus replicons and their IFN-cured counterparts. B. Total cellular RNA was extracted from each cell line and viral RNA quantified by RT-qPCR. Viral RNA copy numbers were normalized to the total input RNA and expressed as genome equivalents/μg of input RNA. The error bars represent the standard deviation determined from four biological replicates. Viral RNA copy numbers were below the limit of detection in both HGT and HGT-Cured cells. The dotted line represents the low limit of detection. C. Detection of viral replication complexes by confocal imaging. Representative merged confocal micrographs showing the detection of dsRNA (green) in indicated cell lines. Nuclei were stained with DAPI (blue). Scale bars correspond to 20μm.

### Cured population of HGT-NV cells demonstrate enhanced HuNoV replication

Stable viral RNA replication under drug-mediated selection leads to an environment where on the one hand host cells face the selective pressure of the drug and, on the other hand, replicating RNA faces innate cellular responses aimed at inhibiting replication. This results in the establishment of a metastable equilibrium prone to adaptation from both cells and the replicating viral RNA. To test for the appearance of such adaptive mutation(s) in the replicon, VPg-linked RNA was extracted from BHK-NV and HGT-NV cells and subjected to consensus genome sequencing. Sequence analysis revealed that viral RNA originating from both BHK-NV and HGT-NV cells underwent many synonymous and non-synonymous genomic changes, suggesting the potential adaptation of HuNoV RNA to specific cellular environments (Supporting information Table 1). To test specifically for host cell adaptation, wild-type VPg-linked RNA extracted from BHK-NV cells was re-transfected in both HGT and HGT-Cured cell lines that were subsequently selected for 5 days with G418. As shown in Fig. 2A, HGT-Cured cells gave rise to a greater number of stable replicon colonies than parental HGT cells. To quantitatively confirm this observation, each cell line was transfected with wild-type VPg-linked RNA replicon and cells were harvested at various time points for RNA extraction and viral RNA quantification (Fig. 2B). From 5 days post transfection, the levels of Norwalk replicon RNA were significantly higher in HGT-Cured cells when compared to HGT cells. Noroviral RNA levels further increased at day 6 to yield a 15-fold difference between HGT-Cured and HGT cells. Quantification of transfected VPg-linked viral RNA at 6h post transfection revealed similar levels in both HGT and HGT-Cured cells suggesting that IFN treatment did not alter the ability of HGT-Cured cells to be transfected (data not shown). We thus concluded that HGT-Cured cells possess a greater degree of permissiveness to viral replication than parental HGT cells.

**Figure 2.**
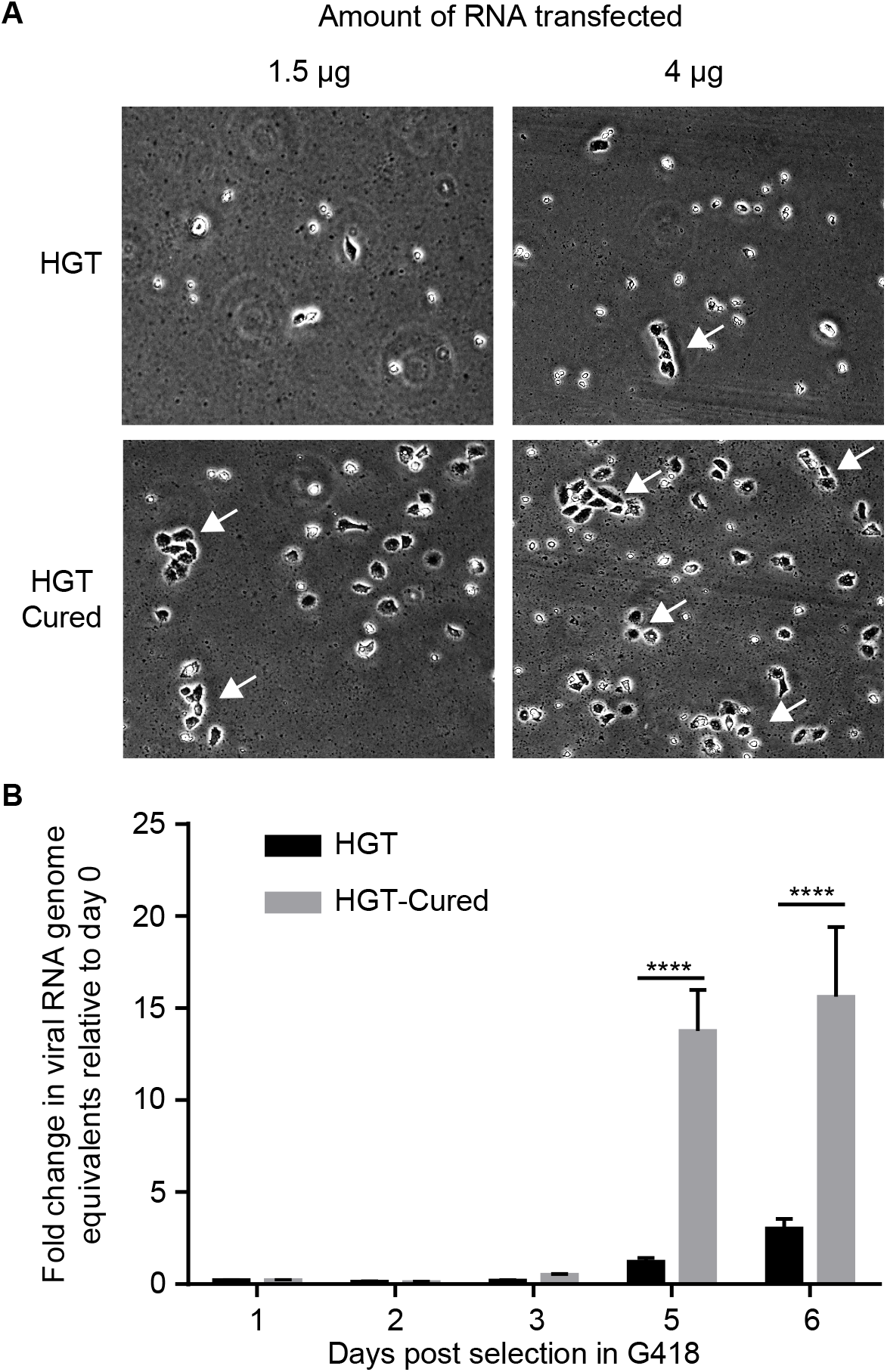
HGT-Cured cells demonstrate enhanced HuNoV replication. A. HGT and HGT-Cured cells were transfected with VPg-linked replicon RNA. After five days of selection with G418 at a concentration of 0.5 mg/mL, cellular morphology was analysed by light microscopy. B. Cells transfected with VPg-linked RNA were harvested at various time points post-transfection for total RNA extraction and viral RNA was quantified by RT-qPCR. Viral RNA levels were determined by comparison to a standard curve and normalized to the RNA input. The data are presented as mean and standard deviation from three replicates. Indicated values are expressed as fold change in genome equivalent normalized to viral RNA levels at 6 hours post-transfection to control for transfection efficiency. Unpaired two-tailed Student’s t-test was used to evaluate the statistical significance.

### An alteration in cellular environment is responsible for enhanced viral replication in HGT-Cured cells

Increased replication of HuNoV VPg-linked RNA in HGT-Cured cells suggests that these cells provide a better cellular environment compared to that of parental HGT cells. To get insights into the mechanism of this cell-derived HuNoV replicon permissiveness, genome-wide expression profiles from HGT, HGT-NV and HGT-Cured cell lines were quantified using Illumina BeadChip microarrays (Tables S2 and S3). We first compared the gene expression profiles of HGT-NV cells relative to their HGT controls. As seen in Figure 3A, more than 2000 genes (HGT-NV-vs-HGT: 2074, HGT-Cured-vs-HGT: 858) were identified for which mRNA expression was significantly modified at a FDR lower than 0.01. Of these, a minority of 151 genes had expression changes in either direction greater than 2-fold suggesting that HuNoV replication has a marginal effect on the whole transcriptional landscape of replicon harbouring cells. Using the same criteria, comparison of gene expression profiles between HGT cells and HGT-Cured cells led to identification of 101 genes for which expression was differentially regulated between the two cell lines (Tables S4-S5). To confirm these observations and to probe for the accuracy of gene expression measured by microarray analysis, we first compared expression changes obtained by microarrays with that measured by RT-qPCR analysis. To this end, ten genes differentially regulated across conditions and spanning a wide range of fold changes were chosen for RT-qPCR validation (Fig. 3B). Globally, we observed a high correlation between the two techniques with Pearson’s correlation coefficients ranging from 0.90 to 0.95. Direct comparison of raw microarray signal intensities with differences in cycle threshold obtained by real-time PCR displayed the same trends suggesting that similar fold changes were primarily the consequence of gene expression differences and were not an artefact resulting from different gene normalization techniques. Differential gene expression at the protein level of selected genes was further confirmed by western blot analysis (Fig. S1A-B). Taken together these data extensively confirms the reliability of gene expression measurements by microarray analysis.

**Figure 3.**
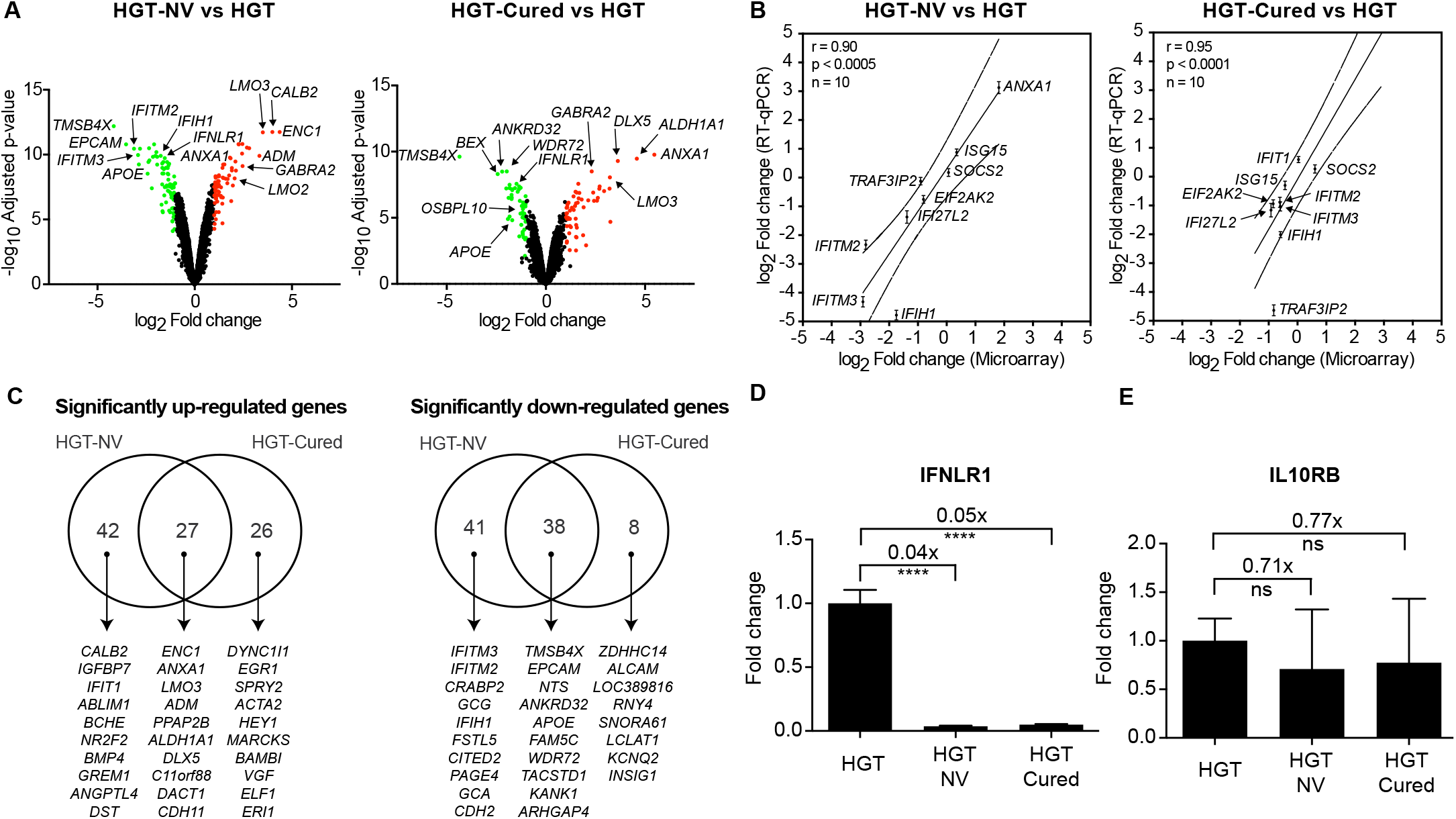
Microarray analysis and validation of differentially regulated genes between HGT-NV, HGT-Cured and parental HGT cells. A. Volcano plots of differentially expressed genes from microarray analysis comparing gene expression in HGT-NV or HGT-Cured compared to parental HGT cells. Significantly up-or down-regulated genes (FDR<0.01 and |log_2_ fold change| ≥ 1) are represented in red or green, respectively. B. Fold change correlation between microarray analysis and quantitative real-time PCR (RT-qPCR). Scatter plots comparing log_2_ fold changes of selected genes measured by microarray analysis and RT-qPCR in HGT-NV or HGT-Cured compared to parental HGT cells. Error bars represent the standard deviation of biological quadruplicate experiments analyzed in triplicate reactions. The Pearson correlation coefficient (r) and the number of pairs analysed (n) are indicated on each graph. Dotted lines illustrate the 95% confidence interval of the linear regression. C. Venn diagrams representing the overlap of significantly up- and down-regulated genes between HGT-NV or HGT-Cured cells compared to parental HGT cells. The top ten genes within each category are shown. D-E. IFNLR1 (D) but not IL10RB (E) gene expression is down-regulated in HGT-NV and HGT-Cured cells. Changes in gene expression of IFNLR1 and IL10RB mRNA levels were normalized to β-actin levels and calculated using the ΔΔ*C_T_* method. Relative expression was determined from one experiment performed in biological triplicate and compared to the expression levels measured in HGT control. Statistical significance was determined using the unpaired *t* test. Error bars represent the standard deviation between biological replicates.

The number of genes showing similar regulation between HGT-NV and HGT-Cured is surprisingly high and likely reflects cellular long-term adaption to replicating RNA (GO term enrichment analysis did not reveal specific pathway enrichment in this gene set). Remarkably, we observed an opposite regulation of several interferon-stimulated genes (ISGs) when comparing the HGT-NV transcriptional landscape with that of parental HGT cells (Fig. 3C and Table S4). While genes coding for IFITM2, IFITM3, IFIH1 and IFI27L2 were down-regulated, IFIT1 and IFIT2 genes were significantly up-regulated in HGT-NV cells. Expression of IFIT1 and IFIT2 proteins is known to be IRF3-dependent (36, 37), while expression of IFITMs and IFIH1 genes was shown to be mediated by ISRE binding (38, 39), this suggests that while HuNoV replication induces IRF3-dependent immune responses, activation of ISRE-dependent genes located downstream in the interferon signalling pathway is inhibited. In line with this observation, microarray analysis identified a statistically significant down-regulation of the type III interferon receptor (IFNLR1) expression in HGT-NV and HGT-Cured cells when both were compared to HGT cells. To confirm and extend this observation, we compared IFNLR1 and IL10RB gene expression by quantitative RT-PCR. Remarkably, we found more than 20-fold decrease in IFNLR1 expression when parental HGT cells were compared to HGT-NV or HGT-Cured cells. In contrast, no significant difference was observed when IL10RB gene expression was measured (Fig. 3D-E).

### Sensing of cytosolic RNA and DNA PAMPs by the innate immune system is functional

To examine whether HGT-NV and HGT-Cured cells were able to detect and mount an innate immune response against cytosolic PAMPs, poly (I:C) and poly (dA:dT) were transfected in the various HGT-derived cell lines and viperin expression levels were measured by RT-qPCR (Fig. 4A-B). Although the cell lines exhibited upregulation of viperin mRNA to various extents, a significant increase of viperin expression above basal levels was detected in all the three cell lines in response to both poly(I:C) and poly(dA:dT). Given that the overall levels of viperin following treatment was comparable across the three cell lines, evident by the relative ratio with respect to β-actin, we concluded that PAMP sensing and downstream signalling pathways are functional in these cell lines.

**Figure 4.**
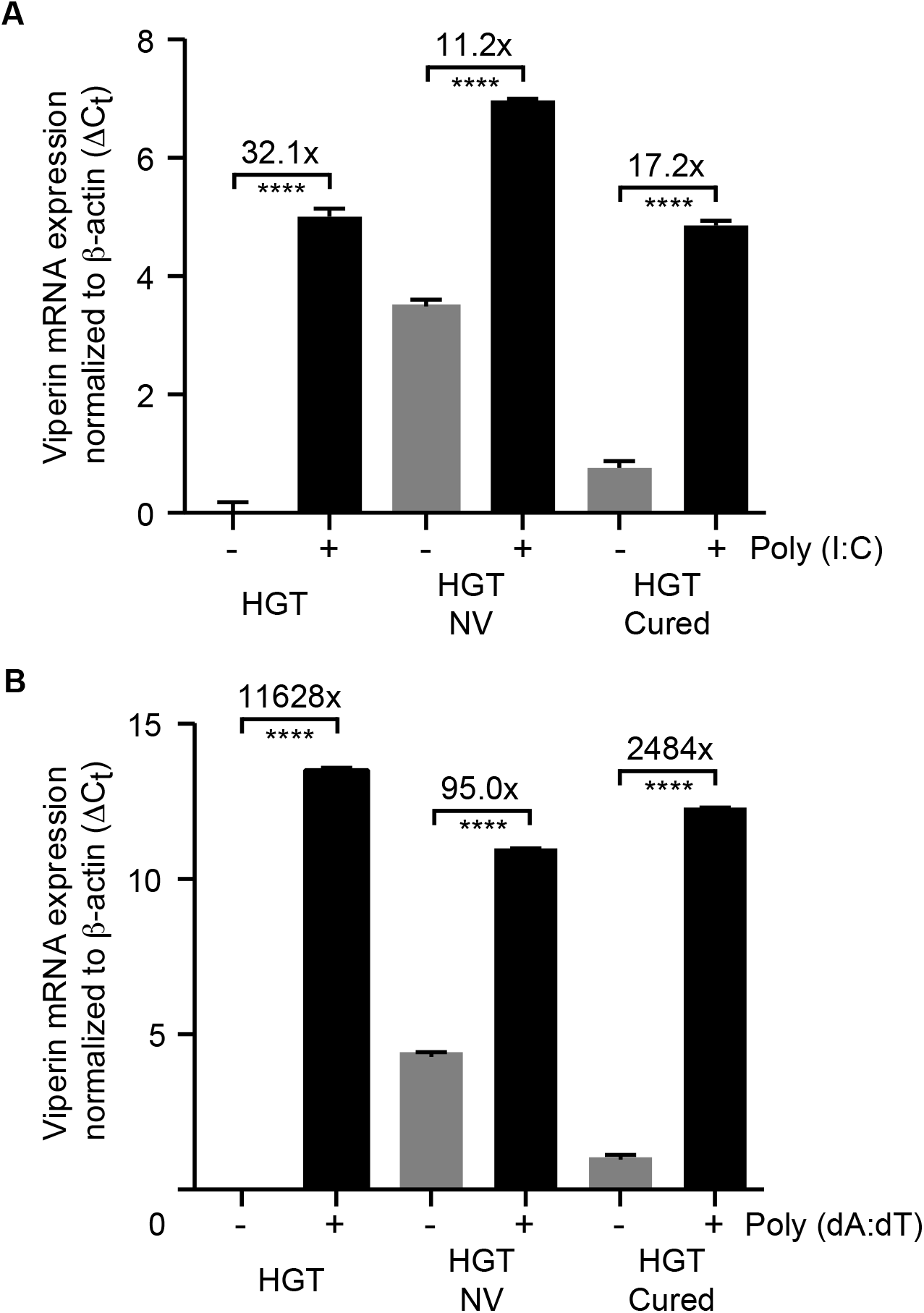
Innate immune responses to RNA and DNA PAMPs are functional. Cells were transfected with poly (I:C) (A) or poly (dA:dT) (B) and total RNA was harvested 8h post-transfection for quantification of viperin mRNA levels by RT-qPCR. Viperin mRNA levels were expressed as differences of cycle threshold (Ct) of viperin, relative to the Ct of beta-actin. The error bars represent standard deviation, determined from the result of three replicates.

### The ability to respond to type III but not type I IFN in HGT-NV and HGT-Cured cells is debilitated

We next investigated the ability of type I and type III interferons to elicit ISGs induction. To this end, cells were incubated with human IFN-β or IFN-λ2 for 16 hours and viperin expression levels were measured by RT-qPCR. We observed that viperin mRNA was readily induced in all three cell lines in response to exogenous type I IFN (IFN-β) treatment (Fig. 5A). However, whereas HGT cells responded to exogenous type III IFN (IFN-λ2) treatment, HGT-NV and HGT-Cured cells did not (Fig. 5B). To test whether the absence of ISG expression is linked to a defect of STAT1 phosphorylation in response to type I or type III interferons, cells were incubated with either recombinant IFN-β or IFN-λ2 and STAT1 phosphorylation status was analysed by western blot using anti-STAT1 phospho-specific antibodies. We observed that although STAT1 was phosphorylated in all cell lines following IFN-β treatment (Fig. 5C), no STAT1 phosphorylation was detected in HGT-NV and HGT-Cured cells following IFN-λ2 treatment (Fig. 5D). To ensure that the absence of STAT1 phosphorylation in HGT-NV cells is not due to a potential clonal effect, different clones as well as a population of the HGT-NV cells were included in the experiment. Similarly, the clones represented here by clone 2 (C2) and the polyclonal cell population responded to IFN-β but not to IFN-λ2 (Fig. 5E-F). Taken together, these results indicate that signal transduction induced by type I IFN is intact while STAT1 phosphorylation induced in response to lambda interferon is inhibited in HGT cells that sustained HuNoV replication.

**Figure 5.**
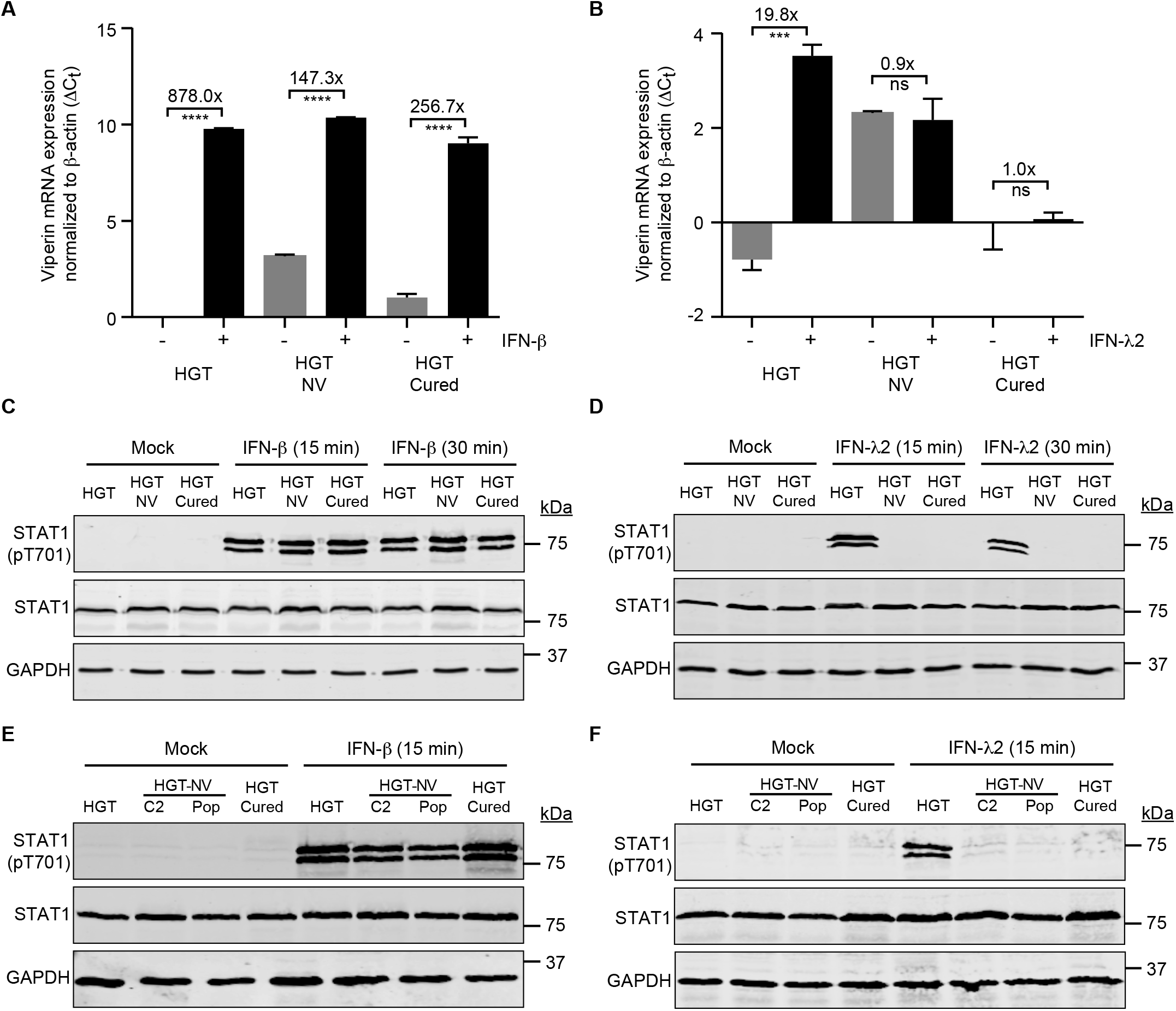
The ability to respond to type III but not type I IFN is debilitated in HGT-NV and HGT-Cured cells. Cells were treated with type I (IFN-β) (A) or type III interferon (IFN-λ2) (B), incubated overnight and harvested for total RNA extraction and quantification of viperin mRNA levels by RT-qPCR. Viperin mRNA expression levels were expressed as differences of cycle threshold (Ct) of viperin, relative to the Ct of beta-actin. The error bars represent the standard deviation, determined from the result of three replicates. (C-F) Western-blot analysis showing STAT1 phosphorylation after stimulation with type I and type III interferons in HGT, HGT-NV and HGT-Cured cell lines (C, E). HGT-NV Clone 2 (C2) and population of cells (pop) were included in the experiment (D, F). GAPDH was used as loading control.

### IFNLR1 overexpression rescues STAT1 phosphorylation in response to IFN lambda

To directly test whether the down-regulation of IFNLR1 expression measured in HGT-NV and Cured cells is responsible for their unresponsiveness to lambda interferon, cells were transfected with an IFNLR1-expressing plasmid and stimulated with IFN-λ2 the day after. We observed that IFNLR1 expression rescued STAT1 phosphorylation in response to interferon lambda suggesting that IFNLR1 down-regulation in HGT-NV and HGT-Cured cells is responsible for IFN-λ insensitivity (Fig. 6). In addition, this experiment shows that the JAK-STAT signalling cascade comprising JAK1, Tyk2 and possibly JAK2 (40) is functional and leads to the phosphorylation of STAT1.

**Figure 6.**
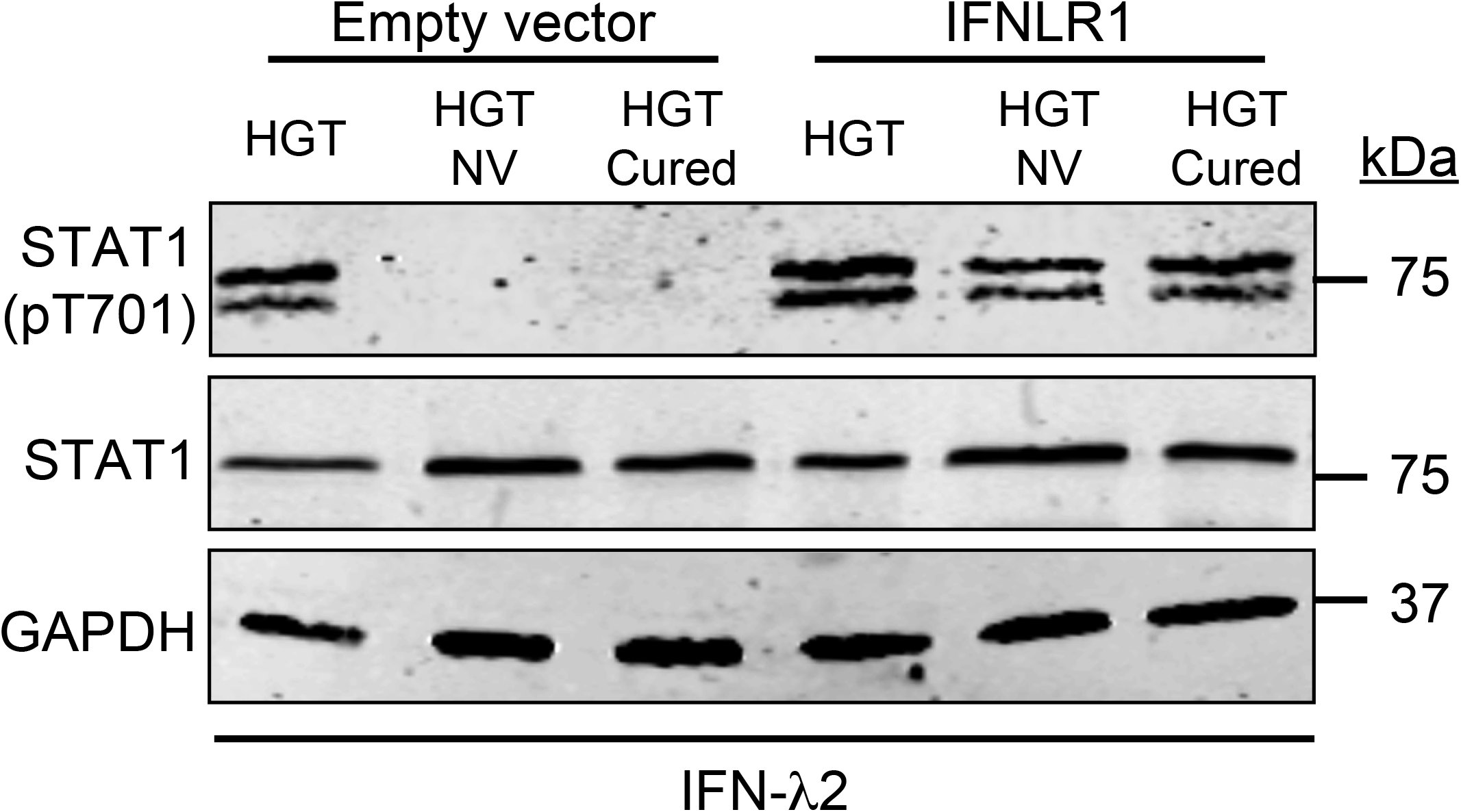
Overexpression of IFNLR1 in HGT-NV and HGT-Cured cells rescues the ability to respond to type III IFN and the loss of the receptor enhances viral RNA replication in HGT cells. HGT, HGT-NV and HGT-Cured were transfected with IFNLR1 expressing plasmid and STAT1 phosphorylation was measured in whole cell lysate by western blot after incubating cells with type III IFN for 15 min (A). STAT1 was used as a positive control and GAPDH was used as the loading control. HGT IFNLR1^−/−^ cells transfected with GI viral VPg-linked RNA were harvested at 0 and 5 days post-transfection for total RNA extraction and viral RNA was quantified by RT-qPCR (B). The viral RNA levels were determined by comparing to a standard curve and normalized to the total input RNA. The data are presented as the mean and standard deviation from at least 11 replicates and are expressed as fold change in genome equivalent at day 0.

### Inactivation of interferon type I and III receptors increases HuNoV replication in epithelial cells

To directly test the influence of type I and type III IFNs on HuNoV replication, HGT cells lines deficient either for the interferon alpha/beta receptor IFNAR1 or for the interferon lambda receptor IFNLR1 were generated using CRISPR/Cas9-mediated genome editing. To this end, parental HGT cells were transduced with lentiviruses expressing IFNAR1 or IFNLR1 single guide RNAs and clonally selected in the presence of puromycin. Individual clones were screened for gene inactivation by IFN-β or IFN-λ2 challenging followed by analysis of STAT1 phosphorylation by western-blot and viperin induction by RT-qPCR (Fig. S2A-B). VPg-linked viral RNA was then transfected and cells were selected in the presence of 0.5mg/mL G418 for up to 6 days. Relative to day 0 taken as reference, we observed a significant increase in genomic viral RNA in all cell lines with the exception of parental HGT cells corroborating the antiviral activity of both type I and type III IFNs on HuNoV replication (Fig. 7). Similarly, an increased ability of modified cell lines to promote colony formation induced by the HuNoV replicon was observed (data not shown).

**Figure 7.**
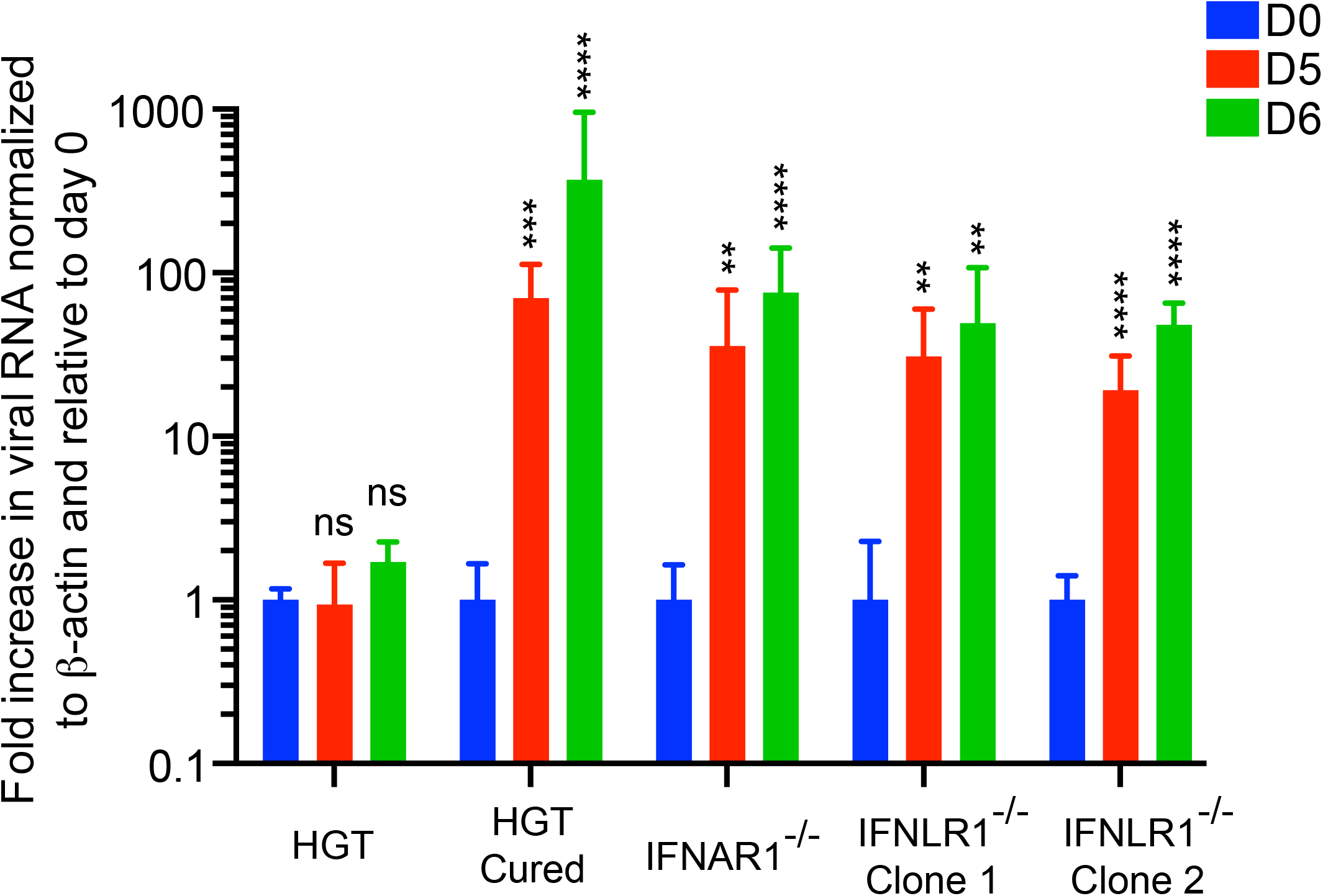
Genetic ablation of IFNAR1 and IFNLR1 interferon receptors promotes HuNoV replication. Parental HGT, IFNAR1 and IFNLR1 knock-out cells were transfected with VPg-linked replicon RNA and submitted to G418 selection. Total cellular RNA was harvested at various time points post transfection and viral RNA was quantified by RT-qPCR. Relative viral replication was determined from one experiment performed in biological triplicate and compared to the replication levels measured in HGT control at day 0. Statistical significance was determined using the unpaired *t* test. Error bars represent the standard deviation between biological replicates.

### IFNLR1 promoter is methylated in HGT-NV and HGT-Cured cells

The dimeric interferon lambda receptor consists of the ubiquitously expressed IL10RB chain subunit and the interferon lambda specific chain IFNLR1 whose expression is limited to cells of epithelial origin (20). Cell-type specific expression of the IFNLR1 subunit was later shown to be inversely correlated with the methylation of its promoter (41). To examine whether viral replication induced, or lead to the selection of, alterations of IFNLR1 gene expression through epigenetic modifications, cells were incubated in the presence 5-aza-2’-deoxycytidine (5azadC), a deoxynucleoside analogue that strongly inhibits DNA methyltransferase (DNMT) activity. As shown in Fig. 8A, we observed a significant 6-fold increase of IFNLR1 mRNA levels in HGT-NV cells when compared to cells incubated with the vehicle only. A statistically significant increase in IFNLR1 gene expression was also observed in the case of HGT-Cured cells but to a lower extent (2.8-fold). In contrast, no change in IL10RB expression was detected when the same cells were incubated with 5azadC (Fig. 8B). In addition, incubation in the presence of MS-275 alone (an HDAC inhibitor) or in combination with 5azadC did not result in increased IFNLR1 expression when compared to 5azadC alone suggesting that the presence of replicating HuNoV did not modify the chromatin structure of the IFNLR1 gene (data not shown). Taken together, these results suggest that the replication of HuNoV in HGT epithelial cells induces a long-term transcriptional silencing of the IFNLR1 gene through the methylation of its promoter.

**Figure 8.**
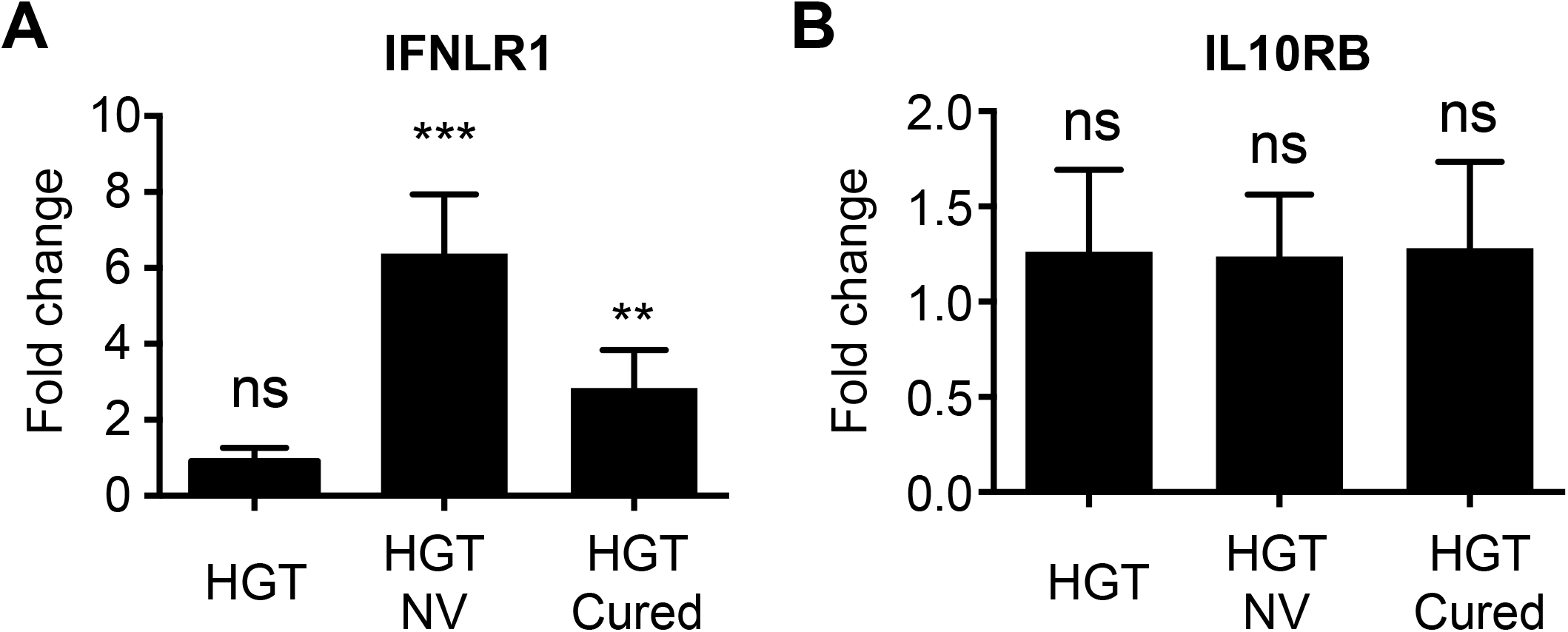
IFNLR1 promoter is methylated during long-term HuNoV replication. HGT, HGT-NV and HGT-Cured cells were incubated in the presence of 10 uM 5azadC for 72 hours or left untreated. Total cell RNA was extracted and IFNLR1 and IL10RB expression was measured by RT-qPCR. Total cellular RNA was harvested 72 hours post-treatment and IFNLR1 and IL10RB mRNA were quantified by RT-qPCR. Relative gene expression was determined from one experiment performed in biological triplicate and compared to the expression levels measured in corresponding untreated cells. Statistical significance was determined using the unpaired t test. Error bars represent the standard deviation between biological replicates.

## Discussion

The purpose of this study was to compare genome-wide transcriptional responses of epithelial cells supporting autonomous HuNoV replication with those elicited by IFN-cured derivatives showing increased viral replication abilities. We found that the interferon lambda receptor is downregulated in cells that sustained HuNoV replication. Mechanistically, this down-regulation of IFNLR1 was mediated by gene methylation as treatment with 5azadC, a DNA methyltransferase inhibitor, rescued the phenotype. This suggests that the lambda IFN pathway is a key constituent of the innate intrinsic defence against human norovirus and shows that an endogenous IFN lambda signalling activity is able to modulate the replication of the virus.

Influence of lambda interferons on murine noroviruses has been highlighted in previous studies using different experimental setups (29, 42, 43). In the case of the human norovirus, recent work using the same replicon in Huh-7 human hepatoma cell line showed that all three types of IFN, when exogenously added, were able to inhibit HuNoV replication leading to virus clearance during long-term treatment (44). However, it is not known whether physiological activation of the interferon pathways, particularly, the lambda IFN signalling pathway exerts an antiviral effect against human noroviruses. The present results add further evidence for the involvement of type III IFN in the control of human noroviruses, but extend earlier findings, by showing that physiological levels of type III IFN signalling can effectively restrict HuNoV replication. One of the major concerns with replicon systems is that gene expression landscapes might reflect clonal selection of RNA-replicating cells rather than a universal impact of viral replication on cellular gene expression. To exclude that differences in gene expression were exclusive to a specific cell type or population, we compared our dataset with previous genome wide-transcription profiles from Huh-7 hepatoma cells supporting autonomous replication of HuNoV RNAs (45). We found a (significant) positive correlation between the two datasets when statistically differentially expressed genes were compared (r=0.19; p=0.03; n=113) (or r=0.49; p=0.1; n=12, FC>2), suggesting that analogous cellular responses were induced in response to HuNoV replication in both cell lines.

It is interesting to note that IFNLR1 is also down-regulated in HGT-NV cells which have not been treated with exogenous type I IFN. This indicates that in the absence of selective pressure mediated by type I IFN, modulation of the type III IFN pathway is preferentially selected over other genes with antiviral properties as exemplified in other studies using RNA replicons with selectable marker. For example, in the case of HCV replicon systems, several antiviral genes including *Viperin* and *MX1* were shown to be silenced through gene methylation, rendering Huh-7 cells permissive to HCV replication (46, 47).

An important observation in this study is the upregulation of several IRF3-dependent genes when HGT-NV cells were compared to parental HGT cells. As IRF3 activation is essential for type I and III interferon induction, upregulation of IRF3-dependent genes suggests that HuNoV replication produces PAMPs that are sensed by RIG-I like receptor(s) which in turn activate the cell-intrinsic immune signalling pathways. Similarly, down-regulation of ISRE-dependent genes in HGT-NV when compared to HGT-Cured cells, which have silenced the IFNLR1 gene to the same extent, put forward the idea that HuNoV replication induces IFN-dependent responses. In line with this, a recent study using the same replicon identified RIG-I and MDA5 proteins as potent negative regulator of HuNoV replication suggesting that viral RNA secondary structures can readily be detected during replication (Dang et al., 2018). These observations are however in striking contrast to the study of Qu and colleagues in which transfection of stool-derived HuNoV RNA readily replicated in 293T cells but failed to induce detectable interferon responses (48). Differences in experimental setups such as including cell lines, virus strains, stable versus transient viral replication and IFN response readouts may account for this discordant observation. Investigating whether diverse strains of HuNoV induce interferon responses in cell-derived human intestinal enteroids will be of interest to probe the influence of HuNoV replication on IFN induction and responses.

Increased replication of HuNoV in HGT-cured compared to either IFNAR and IFNLR1 knock-out cells suggests that differentially regulated genes other than IFN receptors may favour virus replication. However, manual curation of genes differentially regulated between parental and IFN-cured cells did not reveal individual genes convincingly known to modulate viral replication other than IFNLR1 and genes involved in the biosynthesis and trafficking of cholesterol. This suggests either that a low but collective influence of these genes contributes to the increased replication of HuNoV observed in HGT-Cured cells, or that some of the differentially expressed genes identified have a potent yet unknown proviral activity towards the human norovirus.

Overall, our results provide insights into the interactions between the human norovirus and innate cellular responses and show that endogenous levels of λ-IFNs control HuNoV replication suggesting that they may have a therapeutic potential in the treatment of noroviral infections. In addition, the high confidence gene expression datasets provided with this study is expected be useful for the selection and examination of new targets aimed to antiviral therapy.

## Methods

### Cells and Media

Human gastric tumour (HGT) cells, human norovirus replicon-harbouring HGT (HGT-NV) cells and IFN-α interferon-treated HGT-NV cells (HGT-Cured) were maintained in Dulbecco’s minimal essential medium supplemented with 10% fetal calf serum, 2 mM glutamine, 100 U/ml penicillin, 100 μg/ml streptomycin, 1X non-essential amino acid and 0.5 mg/ml G418 in the case of HGT-NV cells. The DNA methyltransferase inhibitor 5-Aza-2’-deoxycytidine (Sigma-Aldrich, A3656) was diluted in sterile water and used at a final concentration of 10 μM.

### Microarray analysis

Four lineages of HGT, HGT-NV and HGT-Cured cells were grown in tissue culture flasks in the appropriate culture media. After four successive passages, total RNA was extracted using TRIzol^®^ (Invitrogen). Microarray analysis was done using the HumanHT-12 v4 Expression BeadChip (Illumina, Chesterford, UK). All microarray experiments, data normalizations and preliminary analysis were fulfilled by the Cambridge Genomic Services, UK.

### Quantitative RT-PCR analysis

Total cell RNA was extracted using a GenElute Mammalian Total RNA Miniprep kit (Sigma) and contaminating genomic DNA was removed through RNase-free DNase I treatment (Roche). Total RNA was then reverse transcribed using random hexamers and the M-MLV RT enzyme (Promega). SYBR green-based quantitative PCR was performed using gene-specific primers listed in (Supporting information Table 6). Each experimental condition was measured in biological triplicate and results are shown as a ratio to levels detected in control cells according to the ΔΔCt method (49). Additional non-template and non-reverse transcriptase samples were analysed as negative controls. Data were collected using a ViiA 7 Real-Time PCR System (Applied Biosystems). Genomic viral RNA was quantified by one step RT-qPCR using GI NV-specific primers. Viral genome copy numbers were calculated by interpolation from a standard curve generated using serial dilutions of viral RNA transcribed from the pNV101 plasmid coding for the full-length Norwalk genome (50).

### Nucleic acid transfection and IFN treatment

Transfections of plasmid DNA or purified viral RNA (NV replicon) were carried out using Lipofectamine 2000 (Invitrogen) according to manufacturer’s instructions. Briefly, cells were seeded in antibiotic-free growth medium at a density of 2×10^5^ or 1×10^6^ cells per well in 24- or 6-well plates, respectively, and incubated overnight at 37°C. Lipofectamine 2000 reagent was diluted in Opti-MEM^®^ (Gibco) and incubated for 5 minutes at 25°C. After the incubation, plasmid DNA or viral RNA, diluted in Opti-MEM^®^ (Gibco), was mixed with the Lipofectamine: Opti-MEM mixture, vortexed briefly and incubated at 25°C for 20 minutes. The DNA or RNA complex was subsequently inoculated onto 80-90% confluent cell monolayers, followed by incubation at 37°C. For viral RNA transfection, media was replaced after 24 h with fresh complete growth medium containing G418 at a concentration of 0.5 mg/ml. For IFN treatments, cells were seeded as described above and incubated at 37°C for indicated time-points with or without recombinant type I IFN (IFN-β; Peprotech, Cat N°:300-02BC), or type III IFN (IFN-λ2; Peprotech, Cat N°:300-02K) at a final concentration of 0.1 ug/ml.

### Immunofluorescence microscopy

Cells were plated on 12 mm glass coverslips and allowed to adhere overnight before fixation with 4% paraformaldehyde in PBS. Cells were then permeabilized for 5 min with PBS-Triton X-100 0.2% and unspecific antigens were blocked for 1h using 2% normal goat serum (Sigma-S2007) in PBS-Tween-20 0.1% (PBST). Cells were then incubated for 1h with primary mouse monoclonal J2 anti-dsRNA antibodies in PBST at a dilution of 1:1000 (J2, SCICONS English & Scientific Consulting, Hungary). After extensive washes with PBST, species-matched AlexaFluor-conjugated secondary antibodies (ThermoFisher Scientific, A-11029) were added at a dilution of 1:500 in PBST for one additional hour. Coverslips were extensively washed and mounted on slides with Mowiol supplemented with DAPI and DABCO. Confocal micrographs were acquired on a Leica TCS SP5 confocal fitted with a 63x 1.3NA oil immersion objective using 405nm and 488nm laser excitation lines under sequential channel scanning to prevent fluorophore bleed-through artefacts due to spectral overlap.

### Western blot analysis

Cell lysates were prepared in radio-immuno precipitation assay buffer (RIPA: 150 mM NaCl, 0.5% sodium deoxycholate, 0.1% SDS, 1mM EDTA, 1% Triton X-100, and 50 mM Tris pH8) supplemented with protease and phosphatase inhibitors (Calbiochem; Cat N°: 539134 and 524625). Protein concentrations were determined by BCA assay (Thermo Fisher Scientific). Equal amounts of total proteins were resolved by SDS-PAGE and transferred to nitrocellulose membranes. Blocking of unspecific antigens was carried out in 5% non-fat dried milk or 5% BSA in PBST for 1 h at 4°C. Primary antibodies were diluted in blocking buffer and incubated overnight at 4°C with gentle rocking (Supporting information Table 7). Membranes were washed three times in PBST for 5 min at RT. Species-matched IRDye-800CW secondary antibodies were diluted in blocking buffer as before and incubated at RT for 1 h. Membranes were washed again three times in PBST for 5 min at RT. Fluorescent signal was detected through an Odyssey CLx infrared imaging system (Li-COR).

### Generation of IFNAR1 or IFNLR1 knockout HGT cells

HGT cells knockout for IFNAR1 or IFNLR1 genes were generated using the CRISPR/Cas9 system. Lentivirus vectors encoding single-guide RNAs against IFNAR1 or IFNLR1 were generously provided by Dr Steeve Boulant and are described in Pervolaraki et al. (51). Vesicular stomatitis virus G-protein-pseudotyped lentiviral particles were generated by transient transfection of 293T cells grown in 6-well plates using 1.25 μg lentiviral vector, 0.63 μg pMDLg/pRRE (Addgene #12251), 0.31 μg pRSV-Rev (Addgene #12253) and 0.38 μg pMD2.G (Addgene #12259) per well. Parental HGT cells were transduced with lentiviral supernatants and incubated for 48 h. Transduced cells were then selected on the basis of their resistance to puromycin at a concentration of 2.5 μg/mL. Clonal isolation was performed by limiting dilution into 96-well plate at a density of 0.3 cell per well and single cell clones were selected on the basis of visual examination. Single cell clones were expanded and tested for IFNAR1 or IFNLR1 gene disruption by RT-qPCR measurement of viperin induction following incubation with IFN-β or IFN-λ2, respectively. Absence of STAT1 phosphorylation following incubation with receptor-matched interferons confirmed the gene ablation.

### Statistical analysis and data sharing

Statistical significance was determined from experiments where n ≥ 3 using two-tailed Student *t* tests in Prism 6.0 (GraphPad). All microarray expression data reported in this study have been deposited into Gene Expression Omnibus (GEO, http://www.ncbi.nlm.nih.gov/geo) with the accession number GSE111041.

## Supplementary figure legends

**Figure S1.** Expression change validation of selected genes at the transcription and translation levels.

A. Direct comparison between microarray signal intensities and difference in cycle thresholds for *ANXA1* and *IFITM3* genes.

B. Validation by western-blot analysis of ANXA1 and IFITM3 protein levels in HGT, HGT-NV and HGT-Cured cell lines. The indicated ratios represent the ANXA1 or IFITM3 protein levels normalized to the corresponding GAPDH loading controls relative to parental HGT cells.

**Figure S2.** Western blot validation of CRISPR/Cas9-induced knockout cells.

A-B. HGT wild-type cells as well as IFNAR1 (A) and IFNLR1 (B) CRISPR/Cas9 clones were treated with type I (IFN-β) (A) or type III interferon (IFN-λ2) (B) for 15 min and STAT1 phosphorylation (pT701) was measured by immunoblot analysis. GAPDH was used as a loading control.

## Author contributions

S.A.E., F.S., M.H., and I.G. designed, performed the research and analysed the data. S.A.E., F.S., and I.G. wrote the manuscript.

## Acknowledgements

We thank Laure Dumoutier (de Duve Institute, Universite Catholique de Louvain) for the gift of the IFNLR1 (LICR2) expressing plasmid. We thank K Green (NIH, Bethesda) and KO Chang for the provision of pNV-Neo.

