## Supplementary figures and images for "Epigenetic suppression of interferon lambda receptor expression leads to enhanced HuNoV replication *in vitro*"

### Supplementary fig 1

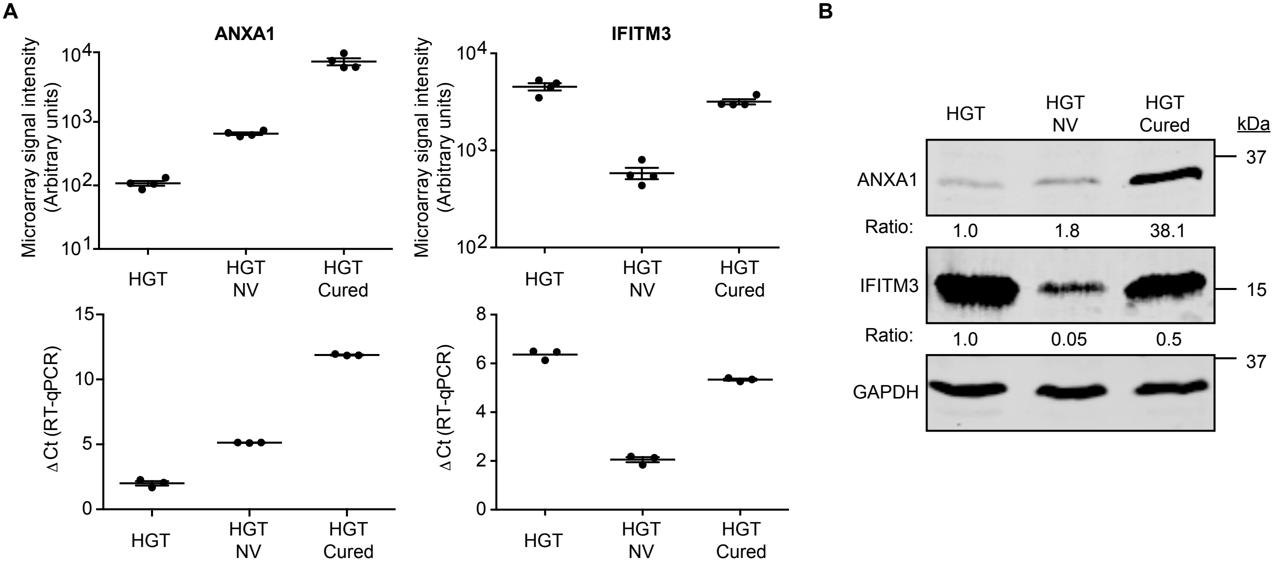

### Supplementary fig 1

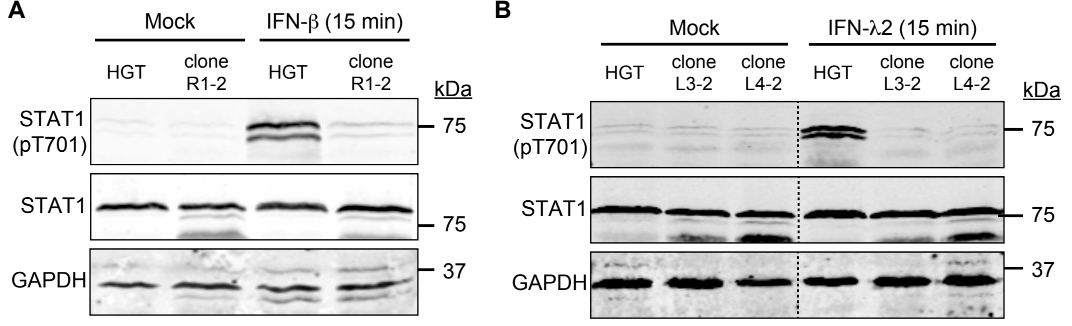
